# Threat expectation does not improve perceptual discrimination despite causing heightened priority processing in the frontoparietal network

**DOI:** 10.1101/2023.07.06.547999

**Authors:** Nadia Haddara, Dobromir Rahnev

## Abstract

Threat cues have been widely shown to elicit increased sensory and attentional neural processing. However, whether this enhanced recruitment leads to measurable behavioral improvements in perception is still in question. Here we adjudicate between two opposing theories: that threat cues do or do not enhance perceptual sensitivity. We created threat stimuli by pairing one direction of motion in a random dot kinematogram with an aversive sound. While in the MRI scanner, 46 subjects (both men and women) completed a cued (threat/safe/neutral) perceptual decision-making task where they indicated the perceived motion direction of each moving dots stimulus. We found strong evidence that threat cues did not increase perceptual sensitivity compared to safe and neutral cues. This lack of improvement in perceptual decision-making ability occurred despite the threat cue resulting in widespread increases in frontoparietal BOLD activity, as well as increased connectivity between the right insula and the frontoparietal network. These results call into question the intuitive claim that expectation automatically enhances our perception of threat, and highlight the role of the frontoparietal network in prioritizing the processing of threat-related environmental cues.

**Significance Statement:** Threatening information receives enhanced priority processing in the brain. Evidence of increased neural activity to threat has fostered the current view that such selective processing leads to a boost in perception, suggesting that motivationally relevant top-down effects can directly change what we see. In the real world, danger is often preceded by an environmental cue that predicts its imminent approach. Here we used an aversive conditioning paradigm to test whether threat cues can change subjects’ ability to visually distinguish between threat and safe stimuli. Our results provide strong evidence for the lack of an effect of threat expectation on perceptual sensitivity, supporting the theory that perception is impenetrable by top-down cognitive influences despite robust neural attentional priority.

## Introduction

Our survival depends on the ability to quickly and accurately identify threats in our environment. Consequently, our brain has evolved to be exceptionally sensitive to danger. Humans detect threat-related stimuli faster (Hansen & Hansen, 1988; Horstmann, 2007; Öhman et al., 2001), make quicker saccades to them (Bradley et al., 2000; Gerdes et al., 2009; Mogg et al., 2007), and look at them more often (Wieser et al., 2009) compared to neutral stimuli. Importantly, this prioritization of threat has been shown to occur not only in fronto-parietal association cortex (Mohanty et al., 2009; Petro et al., 2017) but even as early as the primary visual cortex (Keil et al., 2007; Pourtois et al., 2004; Stolarova et al., 2006), suggesting that it is built into the basic properties of our visual system.

However, while the perception of threat-related stimuli is clearly enhanced, much less is known about the effects of *expecting* a threat. In the real world, dangerous stimuli are often at least partially predictable, such as when we hear a rustling in the bushes before a wild animal attacks or the screeching of tires before a car collision. Therefore, to understand how the brain processes threat, it is critical to understand the mechanisms of brain function when expecting danger.

Two competing theories exist about the effects of threat expectation. On one hand, given that threat stimuli receive enhanced sensory processing, it is natural to assume that expecting threat also enhances perceptual sensitivity (Sussman et al., 2017). On the other hand, perception is often conceptualized as a separate module that cannot be influenced directly by cognition (Firestone & Scholl, 2015), which would indicate that a mere expectation of threat should not enhance perceptual sensitivity.

Surprisingly, while these two hypotheses appear to co-exist in the literature, very little work has been done to adjudicate between them. In the only paper on the subject, Sussman et al. (2017) compared perceptual sensitivity between conditions where subjects were instructed to detect fearful vs. neutral faces. They found enhanced performance for the condition where subjects detected fearful faces and interpreted their results as evidence that threat expectation enhances perception. However, their design did not involve true expectation since neutral and fearful faces were equally likely in both conditions. Instead, it is possible that subjects made more mistakes in the potentially less natural condition of having to detect neutral stimuli (as opposed to the arguably more intuitive condition of detecting a fearful face). Therefore, it remains unknown whether perceptual sensitivity is enhanced in the presence of true expectation, defined as modifying the probability of subsequent stimuli (Summerfield & De Lange, 2014).

Here we adjudicated between the two theories that threat expectation does or does not enhance perception. We employed a fear conditioning paradigm to associate neutral perceptual stimuli (dot motion of different directions) with the likely presence or absence of an aversive outcome (loud noise). We found that expecting a particular type of sensory stimulus strongly biased subjects’ responses toward that stimulus. However, perceptual sensitivity remained constant regardless of whether subjects were led to expect the threat stimulus, the safe stimulus, or were given no expectation at all. Similarly, expecting a threat stimulus had no effect on processing in the visual cortex. Instead, threat cues strongly modulated both the activity and the connectivity profile in frontal-parietal and salience networks, suggesting that threat expectation has strong post-perceptual effects but no direct influence on perception itself.

## Materials & Methods

### Subjects

Forty-nine adult subjects were recruited for the study. We excluded three subjects from analysis due to chance performance (accuracy < 55%), extreme startle reflex, or forgetting to wear contact lenses, respectively. Therefore, a total of 46 subjects (22 females, mean age: 21.7 years, SD: 4.5 years) were included in the analysis. The sample size was determined based on similar fMRI studies on visual perception (Di Luzio et al., 2022; Sussman et al., 2017). Subjects were compensated $20/hour or 1 course credit/hour for a total of 2.5 hours. All subjects were right-handed with normal hearing, normal or corrected-to-normal vision, had no history of neurological disorders, brain trauma, psychiatric illness, or illicit drug use. The study was approved by the Georgia Tech Institutional Review Board. All subjects were screened for MRI safety and provided written informed consent.

### Stimuli and task

Subjects judged the motion direction (left or right) of white dots (density: 2 dots/degree², speed: 5°/s) presented in a black circle (3° radius) in front of a grey background (Fig. 1A). A proportion of dots moved coherently in the right or left direction while the rest of the dots moved randomly. Each dot had a lifetime between three and five frames (refresh rate: 60 Hz) and the coherent motion was carried out by a random subset of dots on each frame. Each dot motion stimulus was preceded by a letter cue (“L” = Left, “R” = Right, “N” = Neutral). The letters L and R predicted the forthcoming stimulus with 75% validity, whereas the letter N was not predictive (both left and right motion were equally likely). We explicitly informed subjects of these contingencies in the instructions. Each trial began with cue presentation for 2, 4, or 6 seconds (chosen randomly), followed by a 3-second dot motion stimulus and an untimed response. A screen with a fixation dot was then presented between trials for a period of 1 or 2 seconds. All stimuli were created in MATLAB using Psychtoolbox 3 (Brainard, 1997; Kleiner et al., 2007; Pelli, 1997).

**Figure 1.**
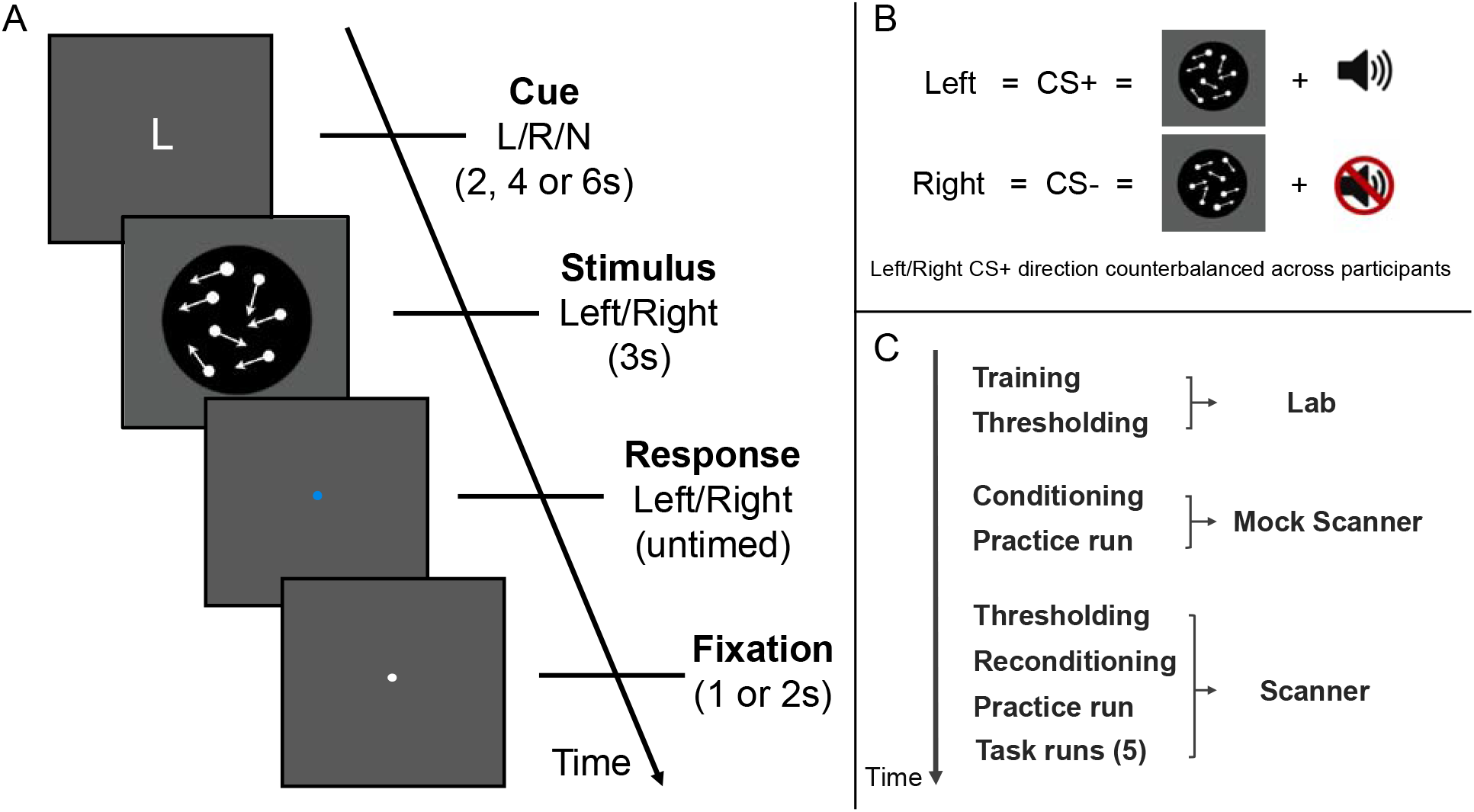
Experimental details. (A) The main perceptual task performed in the scanner. Each trial began with a predictive cue (L = Left, R = Right, N = Neutral) that indicated the likely direction of motion. A random dot kinematogram stimulus was then presented and subjects indicated the direction of motion. (B) The aversive conditioning paradigm. Subjects learned that one direction (left or right, counterbalanced across subjects) was paired with a loud aversive sound delivered through headphones. (C) Depiction of the sequence of events and respective locations of each phase of the experiment.

We used an aversive conditioning paradigm to assign threat (CS+) and safe (CS-) conditions to the otherwise neutral right/left moving dot stimuli (Fig. 1B). The aversive sound consisted of a female scream combined with high-pitched microphone feedback and was delivered through headphones for one second (at 90 dB in the lab and 100 dB in the scanner). The dot motion direction (right/left) that represented the aversive stimulus was counterbalanced across participants. The aversive sound was presented at the end of the dot motion stimulus such that they ended at the same time.

### Experimental design

We split the experiment into three phases that were carried out in the lab, mock scanner, and MRI scanner (Fig. 1C).

### Practice and thresholding outside the scanner

Subjects were first introduced to the task through a 62-trial training outside the MRI scanner in which they learned to perform the task at progressively lower coherence levels. These practice coherence levels were the same for all subjects (50%, 30%, 20%, 12%, 8%, 7%, 6%), starting with a higher coherence of dot motion that gradually decreased, making the task progressively more difficult. Subjects were then familiarized with the three letter cues that indicated whether the motion direction of the upcoming stimulus was most likely to be left (“L”), right (“R”), or neutral (“N”; which gave no information about direction). All subjects received feedback on their accuracy as they went through the practice session. They then completed a 2-up-1-down staircase that determined their individual threshold. The staircase was run until there were 10 reversals and the final coherence level was the average of the last six reversals.

### Conditioning inside mock scanner

After being familiarized with the task, subjects underwent aversive conditioning in a mock scanner that was as similar as possible to the MRI scanner in which they would complete the main task. The conditioning phase consisted of 16 CS+ and 16 CS-interleaved trials (3 seconds each; ISI = 1, 2 or 3 seconds), where 75% of the CS+ stimulus trials co-terminated with a loud aversive sound (i.e., 75% reinforcement rate). We used a high coherence level (30%) during conditioning to establish a clear association with threat and safe directions. Following conditioning, subjects completed a 24-trial practice session (eight trials of each cue) that included the aversive sound. During this practice 50% of the valid CS+ trials co-terminated with the aversive sound. The CS-and Neutral cues were never paired with the aversive sound throughout the experiment.

### Thresholding, reconditioning, and main task inside the scanner

Following the practice in the mock scanner, we prepared subjects for the MRI scanner. In the scanner, they completed the same thresholding task to ensure there were no substantial changes due to the new environment. If there was a difference in lab and scanner thresholds, we used the more conservative (higher) threshold. The motion coherence level (mean = 8.36%, SD = 4.09%) remained constant for all trials of the main task. Before beginning the main task, subjects completed a shorter version of aversive conditioning (24 trials total) to further strengthen the threat/safety association in the scanner environment, as well as a short practice run.

The main experiment included 5 runs each consisting of 4 blocks of 15 trials (60 trials/run; 300 trials total). Subjects were given a 12 second break between blocks and could take longer breaks between runs. In addition to the 60 experimental trials in each run (where no aversive sound was presented), we interleaved 8 extra CS+ valid trials in which the aversive sound was played. We added these extra trials to ensure the conditioned response was not extinguished. These 8 trials were removed from the main analyses.

At the end of each run, subjects reported their level of anxiety via button press (scale of 1-4). They also completed the trait version of the State–Trait Anxiety Inventory at the very beginning of the experiment immediately after the consenting process (Spielberger et al., 1983). These two anxiety measures are outside the scope of the current paper and will be communicated in a separate report.

### fMRI acquisition and preprocessing

We collected the BOLD signal data on a 3T MRI system (Prisma-Fit MRI system; Siemens) using a 32-channel head coil. We acquired anatomical images using T1-weighted sequences (MEMPRAGE sequence, FoV = 256 mm; TR = 2530 ms; TE = 1.69 ms; 176 slices; flip angle = 7°; voxel size = 1.0 x 1.0 x 1.0). We acquired functional images using T2*-weighted gradient echo-planar imaging sequences (FoV = 220 mm; slice thickness = 2.5, TR = 1200 ms; TE = 30 ms; 51 slices; flip angle = 65° voxel size = 2.5 x 2.5 x 2.5, multi band factor = 3, interleaved slices).

We used SPM12 (Wellcome Department of Imaging Neuroscience, London, UK) to analyze the MRI data. Functional images were first converted from DICOM to NIFTI and then preprocessed with the following steps: de-spiking, slice-timing correction, realignment, segmentation, coregistration, and normalization. The functional images were smoothened with a 6 mm full-width-half-maximum (FWHM) Gaussian kernel.

### Behavioral statistical analyses

We computed signal detection theoretic parameters for perceptual sensitivity and response bias (Green & Swets, 1966; Macmillan & Creelman, 2005). We used stimulus sensitivity (d’) to determine subjects’ ability to distinguish between left and right motion directions (i.e., CS+ and CS-). We also calculated response criterion (c) to determine the degree of response bias subjects had for one stimulus type over the other (CS+/CS-). We calculated these measures based on the observed hit rate (HR) and false alarm rate (FAR) such that:

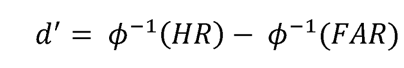

and

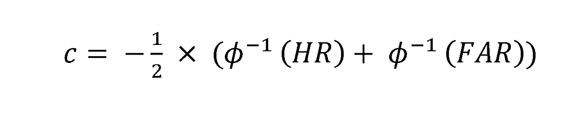

where ϕ^-1^ is the inverse of the cumulative standard normal distribution that transforms HR and FAR into z-scores. HR and FAR were defined by treating the CS+ stimulus as the target. Therefore, negative c values indicate a bias for the CS+ stimulus, whereas positive c values indicate a bias for the CS-stimulus. Additionally, we calculated subjects’ reaction times (RT) for each of the three trial types (CS+ cue, CS-cue, and neutral cue trials). Subjects’ behavioral performance measures (d’, c, RT) during threat expectancy (CS+ cue), safe expectancy (CS-cue) and no expectancy (neutral cue) trials were compared using both Bayesian and frequentist statistics.

### fMRI statistical analyses

We analyzed BOLD signal activity using a general linear model (GLM) in SPM12 (Wellcome Department of Imaging Neuroscience, London, UK). We modeled regressors of interest separately for cue (L, R, N) and stimulus (CS+, CS-), obtained by convolving the unit impulse time series for each condition with the canonical hemodynamic response function. The regressor for each trial included the whole period between the onset of the cue and the offset of the moving dot stimulus. Twelve nuisance regressors related to head motion were included: three regressors related to translation and three regressors related to rotation of the head, as well as their derivatives (Lund et al., 2005). Group-level analyses compared BOLD-signal activity in trials with a CS+ cue versus trials with a CS-or Neutral cue (i.e., CS+ cue trials > CS-& Neutral cue trials). Lastly, we defined the left and right motion areas (MT+) as regions of interest (ROIs) using the MarsBaR toolbox in SPM based on the contrast Stimulus > Fixation (Brett, M., Anton, J. L., Valabregue, R., Poline, 2002). The ROIs were defined as 5 mm spheres centered around peak activity MNI coordinates in right and left MT+ areas separately (Right MT: 46, -68, 0; Left MT: -44, -72, 2). Beta-values were then extracted and compared across cues using an ANOVA.

We further examined whether the functional connectivity between different brain regions was dependent on the threat expectation in a trial (CS+ cue vs. CS-and Neutral cue trials). We used a generalized psychophysiological interaction (gPPI) analysis to determine whether changes in the time-course of whole-brain BOLD activity matches that of a specified seed region in the brain. We conducted the gPPI analysis in the CONN 7 toolbox (Whitfield-Gabrieli & Nieto-Castanon, 2012) using the right insular cortex (coordinates: [47, 14, 0]) – a critical site in the salience network for threat processing (Fullana et al., 2016) – as a seed region.

### Data and code accessibility

All data and codes for the behavioral analyses are freely available at https://osf.io/79x42/. Unthresholded fMRI maps have been uploaded to NeuroVault60 and can be accessed at http://neurovault.org/images/798801/.

## Results

We adjudicated between the hypotheses that threat expectations do or do not enhance perceptual sensitivity. Subjects completed a motion direction discrimination task where one of the two directions of motion was coupled with an aversive stimulus (a loud sound). On each trial, cues indicated whether the threat (CS+) stimulus was more likely, whether the safe (CS-) stimulus was more likely, or whether both stimuli were equally likely. We compared both the behavior and brain activity in the presence of threat expectation (i.e., for trials with a CS+ cue) compared to its absence (i.e., trials with CS-or Neutral cues).

### Effect of CS+ and CS-cues on response bias

As an initial manipulation check, we examined whether subjects took the predictive cues into account by exhibiting a response bias towards choosing the expected stimulus. We quantified response bias using the signal detection theoretic measure for response criterion, c, where negative c values indicate a bias towards the CS+ stimulus (see Methods). We found that the CS+ cue led to a significant bias towards choosing the CS+ stimulus (average c = .30, t(45) = 4.0, p = 2.4 × 10^-4^, Cohen’s d = .59; Fig. 2A), while the CS-cue led to a significant bias towards choosing the CS-stimulus (average c = -.56, t(45) = -8.52, p = 6.0 × 10^-11^, Cohen’s d = -1.26). The criterion values in the CS+ and CS-cue trials were significantly different from each other, and also significantly different from the Neutral cue trials (all p’s < .00015). Therefore, subjects clearly took the cues into account and adjusted their response strategies accordingly.

**Figure 2.**
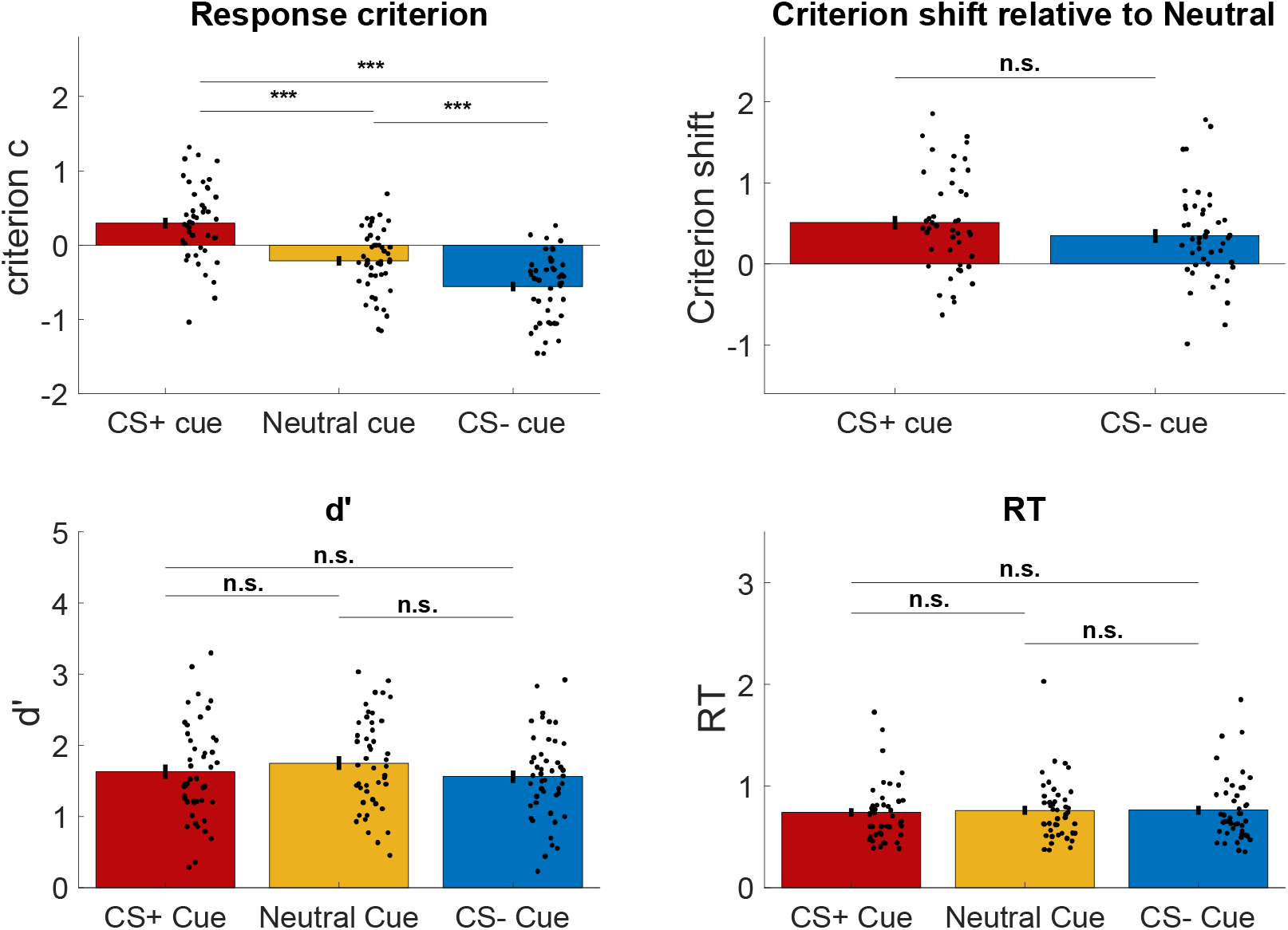
Behavioral effects of threat expectation. (A) Both the CS+ and CS-cues produced a significant response bias towards the corresponding stimulus. Negative c values here indicate a bias toward CS+ responses. (B) Similar criterion shift was observed for CS+ and CS-cues (compared to Neutral cues). (C) Perceptual sensitivity (d’) was similar for CS+ and CS-cues. Sensitivity was slightly higher for Neutral cues presumably due to the noise associated with shifting a criterion from its default value. (D) Reaction time (RT) was similar across all three types of cues. Error bars show SEM, dots indicate individual subjects.

Importantly, we compared the strength of the bias induced by the CS+ and CS-cues. If expecting a threat (but not a safe) cue leads to altered sensory processing, then one may predict that, compared to Neutral cues, there would be a larger response bias shift for the CS+ than the CS-cues. To check for such an effect, we compared the deviation scores c_CS+_ - c_Neutral_ and c_Neutral_ - c_CS-_. We found no significant difference between the two deviation scores (t(45) = 1.15, p = .26, Cohen’s d = .28; Fig. 2B) and a Bayesian t-test suggested that there is substantial evidence for the null hypothesis of no difference (BF01 = 3.4). These results demonstrate that the CS+ and CS-cues had a similar effect on response bias despite the large difference in their affective significance to our subjects.

### Threat expectation does not enhance perceptual sensitivity

Having established that subjects made equivalent adjustments to their response criterion for CS+ and CS-cues, we turned to our main goal of adjudicating between the two hypotheses regarding the influence of threat expectation on perceptual sensitivity (i.e., that threat expectation does or does not improve perceptual sensitivity). To adjudicate between the two hypotheses, we compared the perceptual sensitivity (d’) for each of the CS+, CS-, and neutral cue trials. We found that subjects showed no change in perceptual sensitivity based on cue type (CS+, CS-or Neutral; F(2, 135) = 1.03, p = .36, η_p_ ^2^ = .02; Fig. 2C). A Bayesian ANOVA confirmed that the null hypothesis of no difference was substantially supported (BF01 = 5.9). A post-hoc t-test further established that perceptual sensitivity did not differ between CS+ and CS-cues (t(45) = .83, p = .41, Cohen’s d = .11, BF01 = 4.5). As in previous studies using a cuing paradigm (conducted in the absence of threat expectation) (Bang & Rahnev, 2017; de Lange et al., 2013), the Neutral cues had slightly higher sensitivity than the predictive cues (CS-: t(45) = 2.6, p = .01, Cohen’s d = .30, BF10 = 3.4; CS+: t(45) = 1.45, p = .15, Cohen’s d = .18, BF01 = 2.3), likely due to the noise associated with shifting the response criterion from its default value (Bang et al., 2019; Shekhar & Rahnev, 2021a, 2021b). In addition to perceptual sensitivity, we also examined the effect of cue type on RT and again found no difference between the three cues (F(2, 135) = .08, p = .92, η_p_ ^2^ =.001; BF01 = 13.2; Fig. 2D). Post-hoc t-tests showed no difference between any pair of conditions (all p’s > .3, all BF01 > 4). Therefore, the CS+ cue did not affect either d’ or RT, thus strongly supporting the hypothesis that threat expectation does not enhance perceptual sensitivity.

### Threat expectation does not enhance sensory processing

The behavioral results above strongly suggest that threat expectation does not result in enhanced perceptual discrimination but do not clarify whether threat expectation leads to heightened neural responses in sensory areas. To address this question, we examined the neural activations in the motion sensitive area MT+ as a function of cue type. We defined the left and right MT+ as regions of interests (ROIs; see Methods) and compared the activity in each of the two ROIs for trials with CS+, CS-, and Neutral cues. We found no significant difference in the strength of activation as a function of cue type in either the left MT+ (F(2, 135) = .06, p = .94, η_p_ ^2^ = .001; Fig. 3A) or right MT+ (F(2, 135) = .09, p = .91, η_p_ ^2^ = .001; Fig. 3B). Bayesian ANOVAs showed substantial support for the null hypothesis of no difference between the conditions in both the left (B01 = 13.4) and right MT+ (B01 = 13.09). Post-hoc t-tests produced similar non-significant results (all p’s > .1). These results complement our behavioral findings and suggest that threat expectation not only does not enhance perceptual sensitivity but also does not change neural processing in sensory areas.

**Figure 3.**
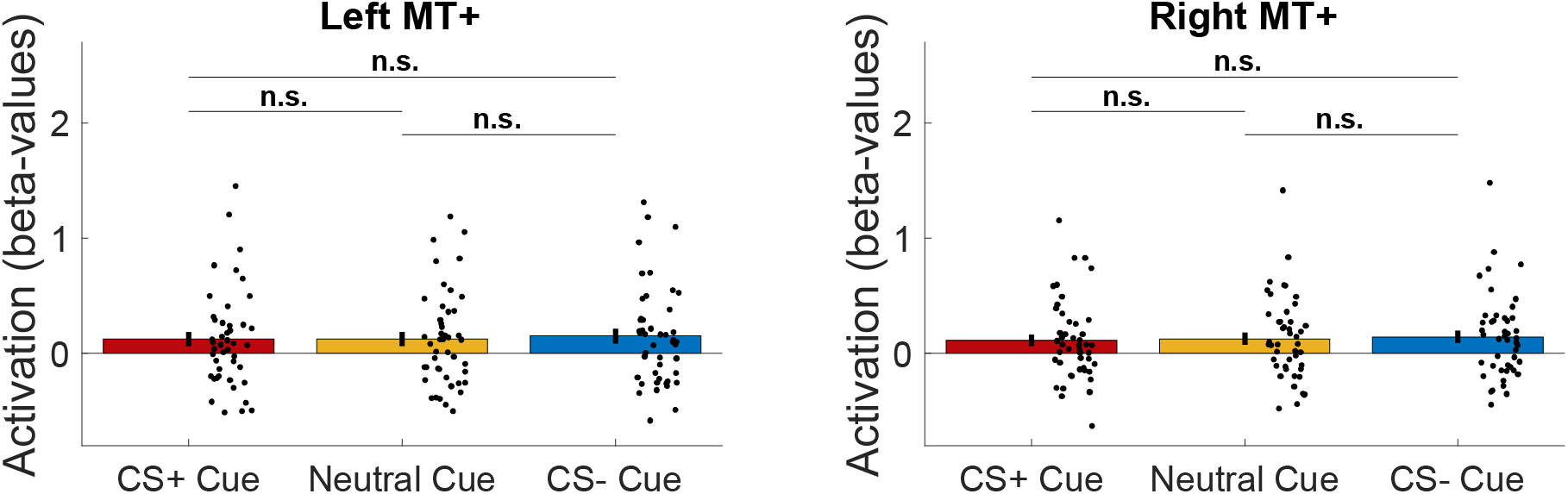
MT+ activations. BOLD signal activations for each cue type (CS+, Neutral, CS-) in both left and right MT+. The three cue types produced equivalent activation in both motion sensitive areas. n.s., not significant.

### Threat cues enhance activity in the frontoparietal and salience networks

Having established that expecting a threat stimulus has no effect on perception or early visual processing, we examined the effects of threat expectation on neural activity in the rest of the brain. Specifically, we performed a whole-brain analysis contrasting CS+ cues to CS-& Neutral cues. We found that threat expectation led to increased activity in regions of both the frontoparietal network (including the right frontal eye field, FEF, bilateral dorsolateral prefrontal cortex, DLPFC, bilateral inferior parietal lobule, IPL, bilateral midcingulate cortex, dorsal precuneus and cerebellum) and the salience network (including bilateral anterior insula and bilateral dorsal anterior cingulate cortex, dACC) (Fig. 4A, Table 1). These results are consistent with the notion that threat cues produce a strong affective response and may indicate high-level prioritizing of threat-related environmental cues (Fullana et al., 2016). On the other hand, the opposite contrast (CS-& Neutral cues > CS+ cues) produced strong activations in somatomotor cortex (Fig. 4B, Table 2), possibly indicating a freezing response for threat cues but not for safe or neutral cues. Similar results were obtained when we compared just the CS+ and CS-cues (Supplementary Figure 1).

**Figure 4.**
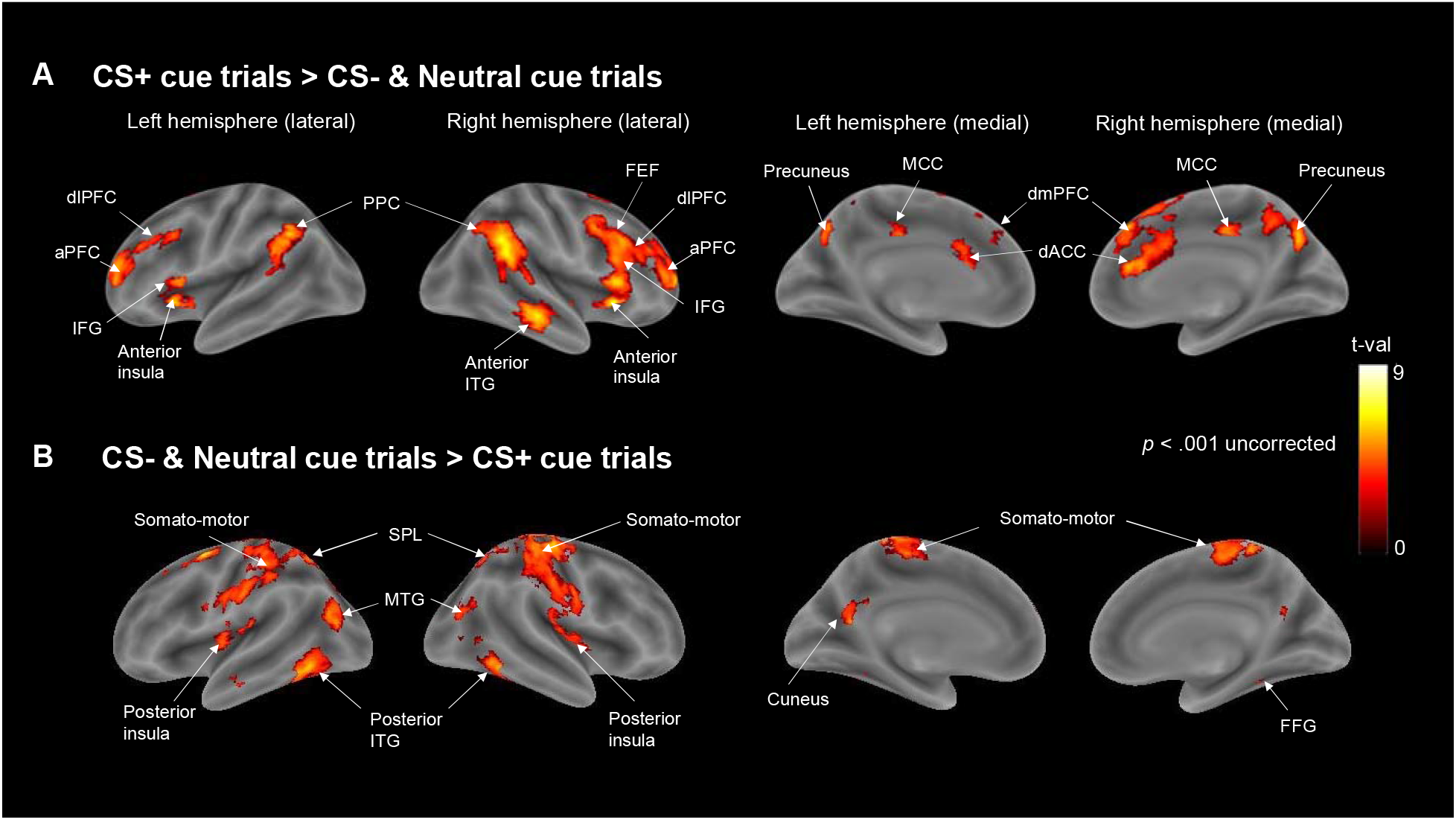
Whole brain comparison between activation produced by CS+ vs. CS-& Neutral cues. (A) Brain activations obtained from the contrast CS+ cue trialslZ>lZCS-& Neutral cue trials. (B) Brain activations obtained from the contrast CS-& Neutral cue trials> CS+ cue trials. The maps were thresholded at p < .001 for display purposes. Colors indicate t-values. aPFC, anterior prefrontal cortex; dACC, dorsal anterior cingulate cortex; dlPFC, dorsolateral prefrontal cortex; dmPFC, dorsomedial prefrontal cortex; FEF, frontal eye fields; FFG, fusiform gyrus; IFG, inferior frontal gyrus; ITG, inferior temporal gyrus; MCC, mid-cingulate cortex; MTG, middle temporal gyrus; PPC, posterior parietal cortex; SPL, superior parietal lobule.

**Table 1.** Whole brain BOLD-signal activation for threat cue trials versus safe and neutral cue trials. Activated clusters for the contrast CS+ cue trials > CS-& Neutral cue trials.

|  | MNI coordinates |  |  |  | Cluster level |  |  |  |
| --- | --- | --- | --- | --- | --- | --- | --- | --- |
|  | Side | x | y | z | Cluster size | t-value | p-uncorr | p-FWE |
| **anterior insula/dIPFC/aPFC** | R | 30 | 24 | -4 | 4411 | 7.72 | 0.000 | 0.000 |
| **Superior medial gyrus** | R | 2 | 38 | 44 | 3024 | 7.71 | 0.000 | 0.000 |
| **Posterior parietal cortex** | R | 60 | -48 | 36 | 1943 | 7.85 | 0.000 | 0.000 |
|  | L | -64 | -48 | 26 | 1102 | 5.71 | 0.000 | 0.000 |
| **Precuneus** | R | 10 | -68 | 34 | 1224 | 6.27 | 0.000 | 0.000 |
| **Middle frontal gyrus** | L | -28 | 52 | 16 | 1186 | 6.04 | 0.000 | 0.000 |
| **Inferior frontal gyrus** | L | -28 | 24 | -4 | 863 | 7.33 | 0.000 | 0.000 |
| **Inferior temporal gyrus** | R | 60 | -26 | -14 | 787 | 6.83 | 0.000 | 0.000 |
| **Cerebellum** | L | -16 | -82 | -32 | 627 | 9.17 | 0.000 | 0.000 |
| **MCC** | R | 6 | -24 | 36 | 170 | 5.46 | 0.002 | 0.017 |
| **Superior frontal gyrus** | L | -20 | 10 | 66 | 127 | 4.89 | 0.005 | 0.051 |

**Table 2.** Whole brain BOLD-signal activation for safe and neutral cue trials versus threat cue trials. Activated clusters for the contrast CS-& Neutral cue trials > CS+ cue trials.

|  | MNI coordinates |  |  |  | Cluster level |  |  |  |
| --- | --- | --- | --- | --- | --- | --- | --- | --- |
|  | Side | x | y | z | Cluster size | t-value | p-uncorr | p-FWE |
| **Somatomotor cortex** | R | 30 | -24 | 64 | 9386 | 6.47 | 0.000 | 0.000 |
| **Inferior temporal gyrus** | R | 54 | -54 | -16 | 1043 | 6.32 | 0.000 | 0.000 |
| **Cerebellum** | L | -48 | -64 | -18 | 880 | 6.48 | 0.000 | 0.000 |
| **Middle occipital gyrus** | L | -46 | -74 | 22 | 548 | 6.00 | 0.000 | 0.000 |
| **Calcarine gyrus** | L | -8 | -58 | 10 | 246 | 5.55 | 0.000 | 0.003 |

**Table 3.**
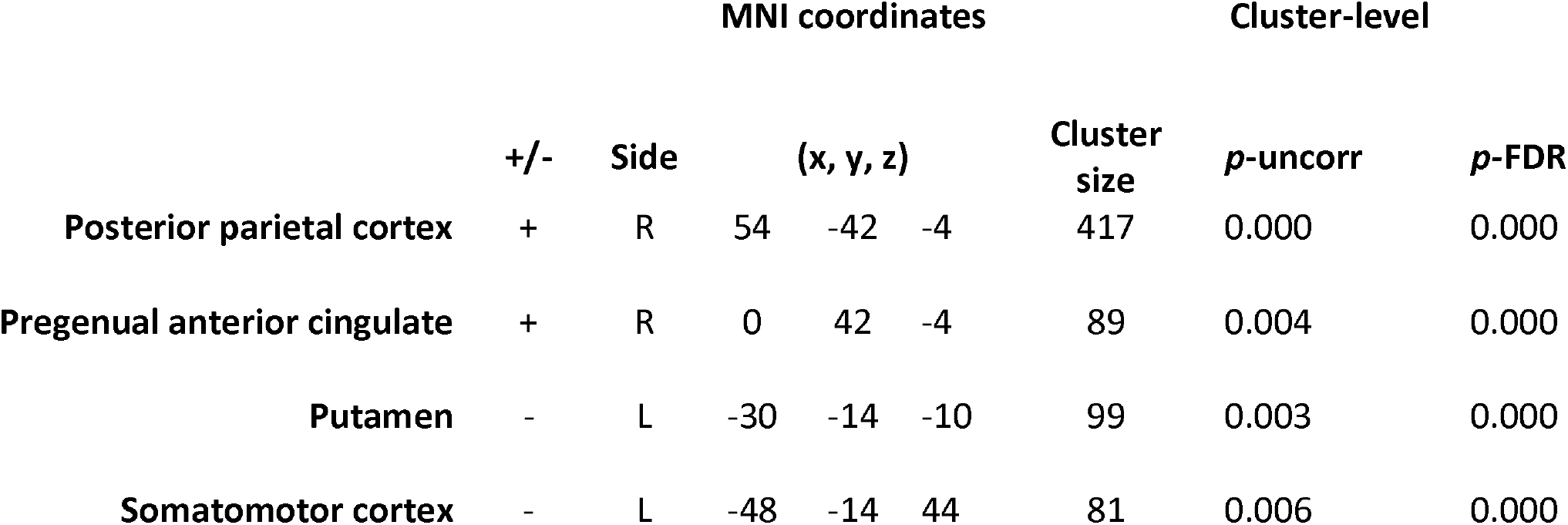
Functional connectivity changes between the right insula and the rest of the brain. Positive values indicate greater effective connectivity during CS+ cue trials, whereas negative values indicate greater effective connectivity during CS-and Neutral cue trials.

|  | MNI coordinates |  |  |  |  | Cluster-level |  |  |
| --- | --- | --- | --- | --- | --- | --- | --- | --- |
|  | +/- | Side | (x, y, z) |  |  | Cluster size | *p*-uncorr | *p*-FDR |
| Posterior parietal cortex | + | R | 54 | -42 | -4 | 417 | 0.000 | 0.000 |
| Pregenual anterior cingulate | + | R | 0 | 42 | -4 | 89 | 0.004 | 0.000 |
| Putamen | - | L | -30 | -14 | -10 | 99 | 0.003 | 0.000 |
| Somatomotor cortex | - | L | -48 | -14 | 44 | 81 | 0.006 | 0.000 |

The interpretation that the increased activity in the frontoparietal and salience networks elicited by the CS+ cues is due to the prioritization of threat-related environmental stimuli leads to the prediction that CS+ cues should also increase the communication between the two networks. Specifically, one may expect increased effective connectivity between areas of the frontoparietal and salience networks in the presences of CS+ cues. To check for such an effect, we compared the strength of the connectivity between the insular cortex (an area known to be highly reactive to threat stimuli, (Fullana et al., 2016)) and the rest of the brain for CS+ vs. CS-& Neutral cues. We found that CS+ cues increased the connectivity between the right insula and both the anterior cingulate gyrus and the right angular gyrus (Fig. 5). Further, consistent with the possibility of a freezing response for CS+ cues, we also found decreased the connectivity between the right insula and motor areas including the somatomotor cortex and left putamen. Overall, these results demonstrate that threat expectation had substantial effect on neural activity in higher levels of processing despite the fact that it did not affect either perceptual sensitivity or early sensory activity.

**Figure 5.**
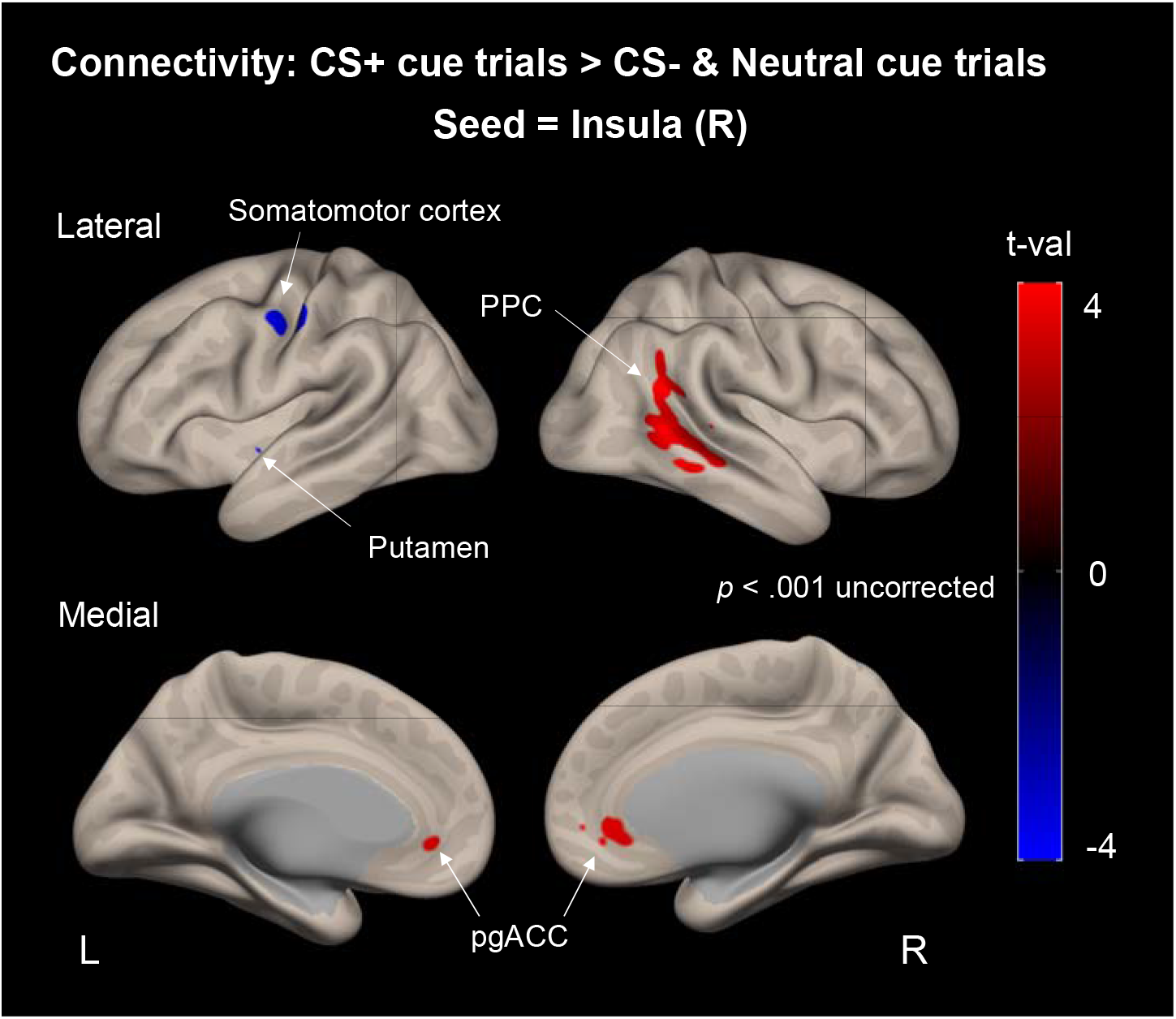
Connectivity differences between CS+ and CS-/Neutral cues. We examined the connectivity differences between the right insular cortex and the rest of the brain. We found that the right insular cortex exhibited increased connectivity with the posterior parietal cortex (PPC) and the pregenual anterior cingulate cortex (pgACC) during trials with CS+ cues. On the other hand, the right insular cortex showed decreased connectivity with the somatomotor cortex and putamen during trials with CS+ cues.

## Discussion

Higher level cognitive states, such as those generated by threatening stimuli, are often thought to penetrate perception and directly change what we see (Pourtois et al., 2013). This view that top-down effects modulate perception and, as a result, change the physical percept is common in studies of threat, emotion, motivation, and beliefs (Firestone & Scholl, 2015). Here we provide evidence that threat information does not enhance nor hinder visual perception. This lack of modulation occurs even though the threat status of an oncoming stimulus is preceded by predictive cues that should, in theory, provide sufficient opportunity for top-down effects to exert their influence. Neurally, we found a lack of modulation of early sensory processing in area MT+ regardless of whether a threat stimulus was expected or not, despite the presence of widespread activations related to attentional priority that were greater for threat-related compared to safe and neutral cues. Taken together, our findings highlight the prioritized neural processing of threat that is consistent with the literature, while showing that such enhanced recruitment does not modulate the early visual areas or change the nature of our visual percepts.

In the real world, dangerous stimuli are often preceded by environmental signals that predict the oncoming threat. Given that many studies have demonstrated enhanced processing and superior performance for threat stimuli (Bradley et al., 2000; Gerdes et al., 2009; Hansen & Hansen, 1988; Horstmann, 2007; Öhman et al., 2001; Stegmann et al., 2020), it appears plausible that the mere expectation of threat stimuli should also lead to better performance. Here we tested this hypothesis using a well-established paradigm where a cue provided probabilistic information regarding the likely identity of an oncoming stimulus (de Lange et al., 2013; Hu & Rahnev, 2019; Kok et al., 2017; Rahnev et al., 2011). We found strong evidence that expecting a threat vs. a safe stimulus had no effect on behavioral performance, as measured both by perceptual sensitivity (d’) and reaction time. Our behavioral results thus indicate that the expectation of threat may not enhance perception.

At first glance, our results seem contrary to a previous study that reported improved perceptual sensitivity to threat compared to neutral cues (Sussman et al., 2017). However, closer examination of the two studies shows that the two findings are not contradictory to each other. In the Sussman et al. (2017) study, subjects discriminated between degraded fearful and neutral faces. In the beginning of each trial, subjects saw a cue that specified the mapping between stimuli and responses. Specifically, the letter F meant that subjects should indicate if the face was fearful or not, such that a “yes” response indicated a fearful face, whereas the letter N meant that subjects should indicate if the face was neutral or not, such that a “yes” response indicated a neutral face. Critically, the cues were not predictive of the actual probabilities of fearful/neutral face presentations (which was always set to 50/50). As such, the cues established a change in stimulus-response mapping by using two different response patterns (fearful faces mapping to “yes” for an F cue but “no” for an N cue) rather than a change in the expectation of the oncoming stimuli. Sussman et al. (2017) found better performance when subjects were detecting fearful compared to neutral faces. However, given that the cues were not predictive (e.g., the letter F was not followed by a fearful face any more than by a neutral face), these findings show the effects of changing the stimulus-response mapping rather than expectation. Specifically, the Sussman et al. (2017) results show that the task set of detecting fearful faces among neutral ones produces better performance, perhaps because it is more intuitive and thus less error-prone, compared to a task set of detecting neutral faces among fearful ones. Our study provided a true expectation cue, such that the CS+ and CS-cues were 75% predictive of the upcoming stimulus and did not use different task sets (i.e., unlike in Sussman et al. (2017), we had a constant stimulus-response mapping between stimuli and responses for the whole experiment). Therefore, the Sussman et al. (2017) results do not contradict our findings here that true stimulus expectation (defined as probabilistic information about the forthcoming stimulus; Summerfield & De Lange, 2014) does not improve perceptual sensitivity.

Our behavioral results were mirrored by a lack of a modulation in early sensory areas responsible for stimulus processing. This was true not only in a whole-brain analyses but even when we defined area MT+ (the sensory area specialized for processing motion stimuli) as a region of interest and tested for differences between the CS+ and CS-cue. Not only did we fail to find a significant difference between the cues, but Bayes Factor analyses showed very strong evidence for the null hypothesis of no difference for both left and right MT+. These results are especially striking in the context of previous studies that have showed clear modulations in sensory areas to the threat stimuli themselves (e.g. Keil et al., 2007; McTeague et al., 2015). These previous studies demonstrate that conditioning paradigms like the current one can lead to significant changes in the early sensory processing. Since early visual areas are capable of threat/safe differentiation (Keil et al., 2007), their lack of activation in our paradigm provides evidence that perceptual sensitivity remains unchanged by the expectation of threat. Our results are in line with established psychophysics findings related to the well-known oblique effect, where vertical and horizontal line orientations are detected faster than oblique orientations, likely due to their prevalence in nature (Appelle, 1972). Despite the consistent increase in perceptual sensitivity for cardinal versus oblique orientations, when high probability predictive cues are introduced there is no enhancement in performance based on stimulus orientation (Stein & Peelen, 2015). Similarly, our data show that even though the visual system is more sensitive to threat stimuli, expecting such stimuli leads to no additional enhancement of perceptual sensitivity.

Critically, the lack of behavioral improvement or changes in sensory areas was coupled with strongly increased activity in the frontoparietal and salience networks. These results complement previous findings that threat stimuli themselves tend to elicit strong activations in these networks (Mohanty et al., 2009; Petro et al., 2017). In our case, these activations are likely to reflect heightened priority processing that is cognitive rather than perceptual in nature.

Overall, we adjudicated between two theories that threat expectation does or does not change perception. We found clear support for the theory that threat expectation does not alter perception, even though it leads to widespread activation in fronto-parietal association cortex. Our results suggest that expectation does not automatically enhance the perception of threat, and highlight the role of the frontoparietal network in prioritizing the processing of threat-related environmental cues.

## Acknowledgements

We thank Ashna Bhardwaj for help with data collection. This work was supported by the National Institutes of Health (Grant R01MH119189) and the Office of Naval Research (Grant N00014-20-1-2622).

