## Supplemental Fig. 1 for "Threat expectation does not improve perceptual discrimination despite causing heightened priority processing in the frontoparietal network"

**Supplementary**

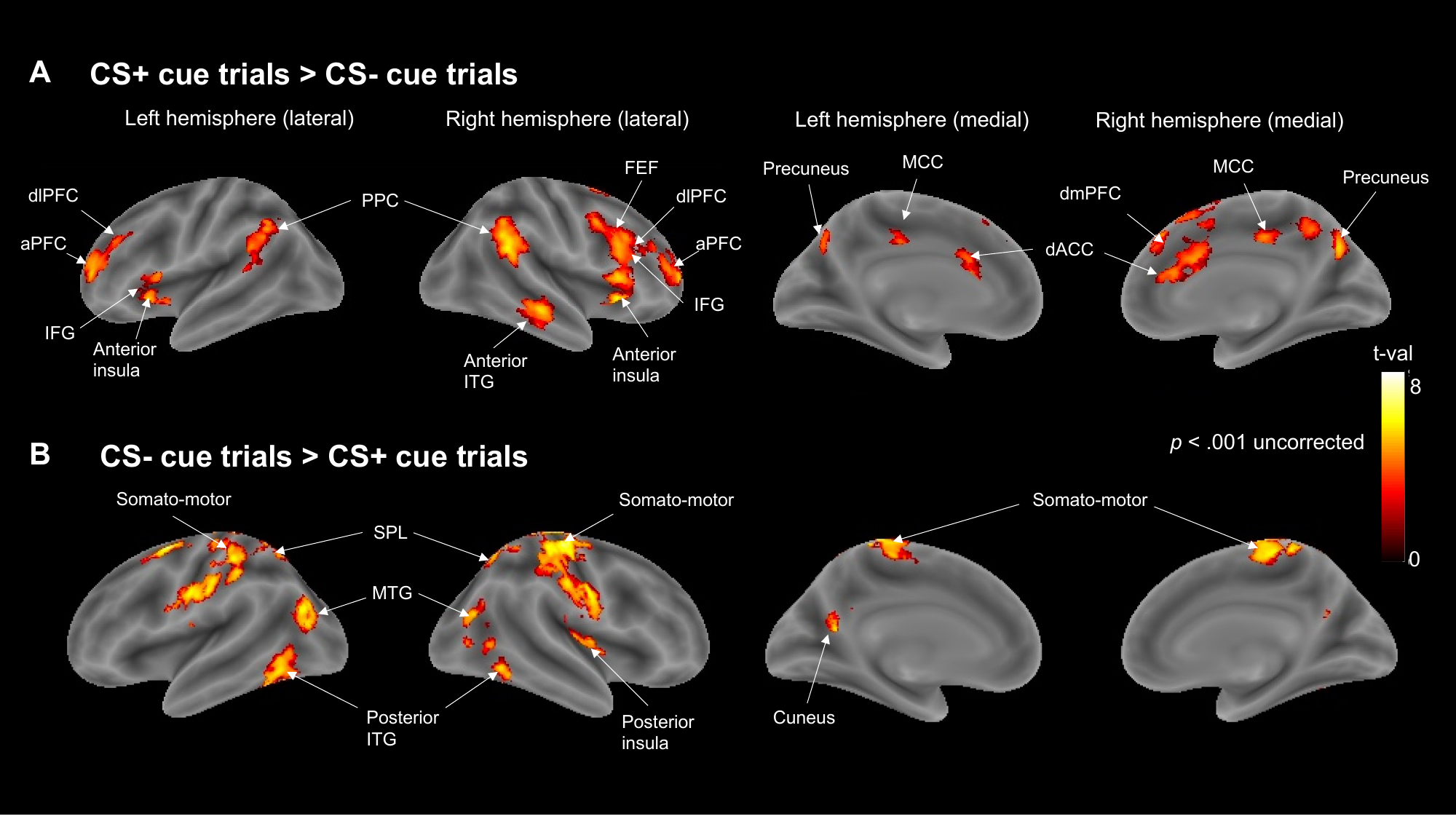

**Supplementary Figure 1.** Whole brain comparison between activation produced by CS+ cue trials vs. CS- cue trials. (A) Brain activations obtained from the contrast CS+ cue trials > CS- cue trials. (B) Brain activations obtained from the contrast CS- cue trials > CS+ cue trials. Activation is very similar to CS+ cue trials > CS- & Neutral cue trials and vice versa. Heat map indicates t-values at *p* > .001. Regions depicted: Anterior inferior temporal gyrus; Anterior insula; aPFC, anterior prefrontal cortex; Cuneus; dACC, dorsal anterior cingulate cortex; dlPFC, dorsolateral prefrontal cortex; dmPFC, dorsomedial prefrontal cortex; FEF, Frontal eye-fields; IFG, Inferior frontal gyrus; MCC, mid-cingulate cortex; MTG, Middle temporal gyrus; Posterior inferior temporal gyrus; PPC, Posterior Parietal Cortex; Precuneus; Somatomotor/Somatosensory cortex; SPL, Superior parietal lobule.
